# Revisiting the Humidity Ramp Protocol for Assessing Human Heat Tolerance Limits

**DOI:** 10.1101/2025.11.09.686670

**Authors:** Fèlix Faming Wang, Yi Xu, Haojian Wang, Min Cui, Xue Hou, Boan Wei, Xiong Shen

## Abstract

**Background:** Humidity ramp protocols are widely used to estimate human heat tolerance limits, yet reported critical environmental limits (CELs) vary markedly across studies. Whether this variability reflects true physiological differences or systematic methodological artifacts related to ramp design remains unresolved.

**Methods:** We combined first-order thermal modeling with controlled physiological trials to examine how ramp temporal structure (step dwell time) influences apparent rectal temperature (*T*_*cr*_) inflection points. Twenty-six healthy young adults completed randomized trials at 42 °C under two protocols: an *aggressive-ramp* (30-min equilibration followed by humidity increases every 5 min) and a *slow-ramp* (4-h equilibration followed by hourly humidity increments). CELs derived from ramp protocols were further evaluated using prolonged fixed-condition exposures in independent cohorts (*n*=14 per sex).

**Results:** Short-step dwell times (Δ*t*/*τ* ≪1) prevented rectal temperature from approaching its step-specific equilibrium, resulting in kinetically constrained, non-equilibrium dynamics and earlier *T*_*cr*_ inflection points. Consequently, aggressive-ramp protocols yielded substantially lower CELs than slow-ramp protocols in both sexes (≈3.5 °C difference). CELs derived from the slow ramp closely matched those obtained from prolonged fixed-condition exposures, whereas aggressive ramps misclassified physiologically compensable conditions as uncompensable.

**Conclusion:** Rapid humidity increments systematically underestimate heat tolerance because environmental forcing outpaces physiological response kinetics. Accurate CEL determination requires either prolonged fixed-condition exposures (benchmark approach) or sufficiently slow ramps that allow near-equilibrium responses (Δ*t*/*τ* ≳0.3–0.5).

## 1. Introduction

As global temperatures rise and extreme heatwaves become more frequent, intense, and prolonged (1), defining the physiological limits of human heat tolerance has become increasingly urgent (2–5). A central challenge is determining the critical environmental limits (CELs) beyond which the body can no longer maintain heat balance. CELs are commonly expressed using wet-bulb temperature (*T*_w_), which integrates dry-bulb temperature (*T*_db_) and humidity to quantify heat stress. Exceeding these limits leads to uncompensable heat stress, characterized by sustained increases in rectal temperature (*T*_cr_) and elevated risk of heat-related illness (6, 7).

A widely used approach to identify CELs is the *humidity ramp protocol* (also termed the *thermal-step* or *inflection-point* method), originally developed by Belding and Kamon (8) as a practical tool to estimate safe exposure durations and evaporative coefficients for prolonged occupational work in hot conditions. This method incrementally increases humidity in fixed steps (typically 1 mm Hg every 10 min) following an initial stabilization period (~60 min) at constant *T*_db_ (9). In practice, however, chamber humidity reaches each new setpoint within minutes (2–5 min), such that these “ramps” are implemented as a sequence of discrete step changes rather than truly continuous transitions. The onset of a sustained rise in *T*_*cr*_ is interpreted as the transition from compensable to uncompensable heat stress. Although conceptually straightforward, implementations vary considerably in stabilization duration, step magnitude, and ramp rate (8–16). For example, some studies employ short stabilization periods (~30 min) followed by rapid humidity increments every 5 min (e.g., 1 mm Hg increases in water vapor pressure or 3% relative humidity increments; [10–14]), whereas others adopt longer equilibration phases (70-min; [15, 16]) and slower stepwise increases (e.g., 1 mm Hg water vapor pressure increments every 10 min; [8, 9]).

Despite broadly similar participant characteristics, reported CELs vary substantially, ranging from ~26 °C to 33 °C in young adults under comparable metabolic conditions (1.3–1.5 Mets) (10–16). Such variability is difficult to reconcile with physiological differences alone and raises a fundamental methodological question: does the temporal structure of ramp protocols — particularly step dwell time— systematically bias CEL estimation?

Recent work from our group demonstrated that environments classified as uncompensable using humidity-ramp protocols remained fully compensable under prolonged fixed-condition exposures, the gold standard for assessing environmental compensability (17). This discrepancy raises a critical question: do *T*_cr_ inflection points observed during ramp protocols reflect true physiological limits, or are they shaped by the temporal dynamics of the protocol itself?

A key but often implicit assumption of ramp protocols is that the thermoregulatory system approaches near-equilibrium conditions following each environmental step. However, the short dwell times commonly used (5–10 min) are unlikely to permit full expression of thermoregulatory heat loss responses (17). When environmental change occurs on time scales shorter than the physiological response, environmental forcing outpaces thermoregulatory adjustment, and the system operates under kinetically constrained, non-equilibrium conditions. Under such conditions, core temperature does not reach its step-specific equilibrium but instead reflects a transient trajectory governed by incomplete adjustment. Importantly, this state is fundamentally distinct from true uncompensable heat stress (18– 20), which arises only when the required heat loss exceeds the maximal achievable heat dissipation (21, 22). In this context, the observed *T*_cr_ inflection may represent a premature deviation from heat balance rather than a true physiological limit.

To test this hypothesis, we examined whether apparent CELs are governed by ramp dwell time. We combined controlled human experiments with first-order thermal modeling to characterize *T*_*cr*_ responses across distinct ramp structures while minimizing inter-individual variability. We hypothesized that short dwell times induce kinetically constrained, non-equilibrium dynamics, leading to earlier inflection points and systematic underestimation of CELs. Together, our findings identify a fundamental source of bias in humidity ramp protocols and provide a mechanistic framework for improving their physiological validity.

## 2 Methods

### 2.1 Model derivation and assumptions

The temporal evolution of rectal temperature (*T*_*cr*_) during progressive humidity changes was derived from the first principles of whole-body heat balance. The human body is modeled as a single, lumped thermal compartment that exchanges heat with the surrounding environment via convection, radiation, and evaporation. At any time *t*, the rate of body heat storage equals the rate of metabolic heat production minus the combined dry (convective and radiative) and evaporative heat losses (23)

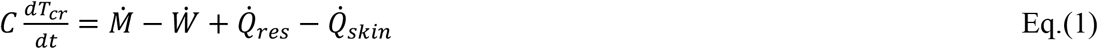

where, *C* is the effective whole-body heat capacity 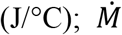 is metabolic heat production rate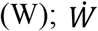is external mechanical work rate (Watt; 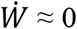 watt under seated conditions); 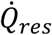 and 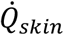 are respiratory and skin heat exchange rates (W), respectively.

The net heat loss through the skin surface can be expressed as the sum of convective, radiative, and evaporative fluxes (23)

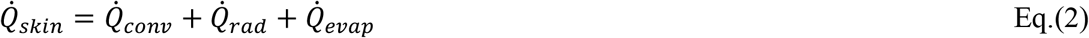

with 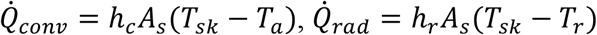 and 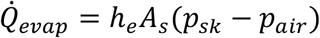
where, *h*_*c*_, *h*_*r*_, and *h*_*e*_ are the convective, radiative, and evaporative heat transfer coefficients (W/m^2^/°C), respectively; *A*_*s*_is body surface area (m^2^); *T*_*sk*_ is mean skin temperature (°C); *T*_*a*_ and *T*_*r*_ are ambient air (dry-bulb) and mean radiant temperatures(°C), respectively; *p*_*sk*_and *p*_*air*_are water vapor pressures at the skin surface and in the ambient air, respectively (Pa).

Under the classical humidity ramp experimental conditions — where air temperature is held constant and relative humidity progressively increases — metabolic heat production 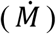 and respiratory heat exchange 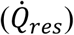 can be treated as approximately constant. Consequently, variations in total heat loss are primarily governed by humidity-dependent changes in the potential for sweat evaporation (24, 25). To obtain a tractable representation of the system’s transient thermal response, the net heat exchange was linearized around the steady-state operating point:

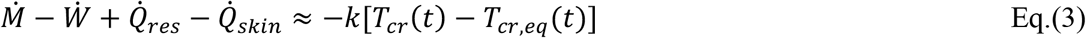

where, *T*_*cr,eq*_ is the equilibrium rectal temperature (i.e., the asymptotic steady-state value under fixed environmental conditions) that would be achieved if the environment were held constant indefinitely (°C). *k* is an effective whole-body heat transfer coefficient (W/°C) — the lumped sum of convective, radiative, and evaporative heat conductances, modulated by physiological control of skin blood flow and sweating. Mathematically, *k* represents how efficiently excess body heat is dissipated when *T*_*cr*_exceeds equilibrium, which is read as:

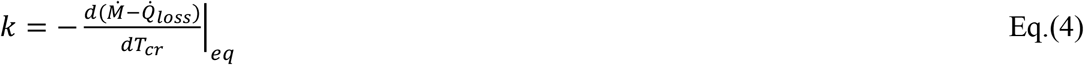

where, 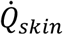 denotes the sum of convective, radiative, evaporative, and respiratory heat losses, i.e.,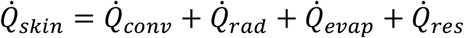

Substituting Eq.(3) into the whole-body energy balance [i.e., the Eq.(1)] yields

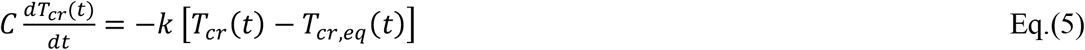

This relation indicates that the rate of rectal temperature change is proportional to the instantaneous deviation from the environment-defined equilibrium value, reflecting the finite response time of the thermoregulatory system to environmental perturbations.

Dividing both sides of the Eq.(5) by *C* and defining the time constant 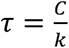 gives

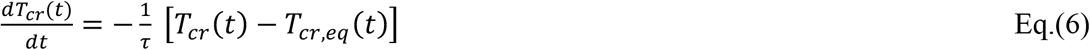

The thermal time constant τcharacterizes the rate at which *T*_*cr*_ approaches equilibrium following an environmental step change. Eq.(6) describes a linear first-order low-pass response with time constant *τ* that encompasses both passive body heat storage and active thermoregulatory feedback. The time constant *τ* represents the characteristic response time for *T*_*cr*_to approach a new equilibrium following an environmental step change in relative humidity (RH). This formulation captures the first-order response dynamics of the thermoregulatory system under time-varying environmental conditions.

During each discrete humidity step of duration Δ*t*, the equilibrium temperature *T*_*cr,eq*_is assumed to remain constant. The analytic solution of Eq.(6) for a step initiated at time *t* = *t*_*n*_ is given by

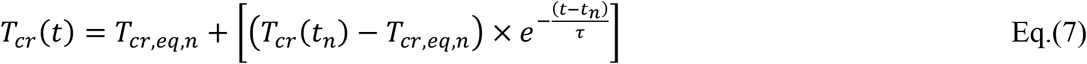

At the end of each step (*t*_*n+1*_ = *t*_*n*_ + Δ*t*):

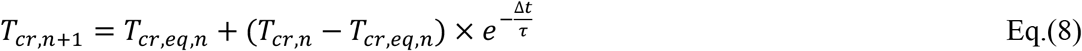

Eq.(8) quantifies how far the body approaches equilibrium within each step. If Δ*t* is sufficiently small, Eq.(8) can be approximated using the Euler form (26)

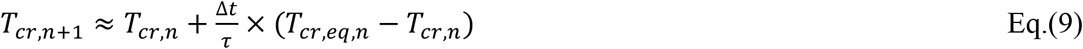

When environmental humidity changes occur on timescales shorter than the physiological response time (i.e., Δ*t*/*τ* ≪1), the thermoregulatory system cannot approach its step-specific equilibrium within each increment, leading to the accumulation of residual deviations from equilibrium across successive increments. As expressed by

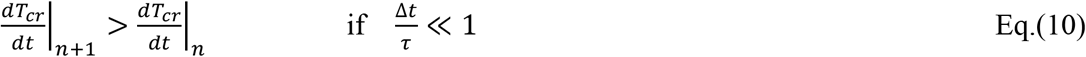

where, 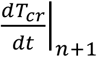 and 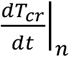 denote the slopes of rectal temperature change at the (*n*+1)th and *n*th steps, respectively. This progressive increase in slope leads to a premature, apparent inflection point in *T*_*cr*_.

Importantly, this behavior arises from kinetically constrained, non-equilibrium dynamics, rather than indicating a true transition to uncompensable heat stress, which requires sustained imbalance under steady or quasi-steady conditions. Detailed mathematical justification is provided in the Supporting Information.

### 2.2 Physiological verification study

#### 2.2.1 Sample size and power calculation

The primary comparison focused on differences in the critical inflection point of rectal temperature (*T*_*c*r_) between two ramp protocols (aggressive- and slow-ramp) and between sexes. Power analysis was performed based on expected sex differences, assuming a large expected effect size (Cohen’s *d* = 0.8) and a paired-samples *t*-test. This indicated that a minimum of 12 males and 12 females would provide 80% power at a two-tailed significance level of 0.05.

#### 2.2.2 Participants and ethical approval

Fourteen healthy young males (age: 23.6±2.0 yrs, height: 1.77±0.07 m, weight: 74.9±9.2 kg, body surface area: 1.92±0.15 m^2^, body mass index: 23.9±1.5 kg/m^2^) and twelve healthy young females (age: 22.5±2.1 yrs, height: 1.62±0.07 m, weight: 55.3±6.1 kg, body surface area: 1.58±0.10 m^2^, body mass index: 21.1±2.4 kg/m^2^) participated. All were non-smokers and free from cardiovascular, metabolic, or thermoregulatory disorders.

Female participants were tested during the follicular phase (days 1–12 following menses onset), when progesterone levels are low, and none used hormonal contraceptives. Written informed consent was obtained prior to participation. The study was approved by the Institutional Review Board (IRB) of xxx (information removed due to double blind requirements) and conducted in accordance with the Declaration of Helsinki.

#### 2.2.3 Experimental design and rationale

Upon arrival, hydration status was assessed via urine specific gravity (USG), with testing initiated only when USG ≤1.020 (27). Participants exceeding this threshold consumed a ~37 °C isotonic electrolyte solution until euhydration was achieved. Baseline thermal state was standardized by requiring a thermoneutral rating and an initial *T*_*cr*_≤ 37.1 °C. To maintain stable environmental conditions throughout each trial, a restroom facility was installed within the chamber, allowing participants to remain inside continuously and thereby avoiding disturbances associated with chamber exit and re-entry. Wet-bulb temperature (*T*_*w*_) was calculated using the Stull equation (28).

Each participant completed two physiological trials (aggressive-ramp and slow-ramp) in a randomized, counterbalanced order, separated by at least one week. Experiments were conducted in a convective heat chamber maintained at *T*_db_ = 42 °C with negligible radiant heat load. Participants remained seated and performed light activities (e.g., reading, typing, or using mobile devices). In the aggressive-ramp protocol, relative humidity (RH) was set at 28% for 30 minutes, followed by 2% increases every 5 minutes until termination. In the slow-ramp protocol, participants underwent a 4-hour equilibration at 40% RH, followed by stepwise increases of 6% RH per hour from hours 4–6, and 3% RH per hour thereafter.

The slow-ramp protocol was initiated at 40% RH to ensure experimental feasibility while limiting prolonged exposure-related confounders (e.g., fatigue and circadian influences). Initiating at lower RH (e.g., 28% in aggressive-ramp) would have substantially extended exposure duration before reaching uncompensable conditions. Importantly, even when RH values overlap between protocols, the effective thermal load differs due to large differences in dwell duration (5 min vs 60 min), resulting in distinct physiological states.

Although these protocols are often described in terms of nominal ramp rates (e.g., %RH per hour or per 5 minutes), each humidity adjustment was implemented as a discrete step. In practice, the chamber reached each new target RH within approximately 2–3 minutes, after which conditions remained stable for the remainder of the dwell period. Participants were therefore exposed to a sequence of stepwise environmental transitions rather than a continuous ramp. Under these conditions, the primary determinant of thermophysiological responses is the dwell duration (Δ*t*) at each step relative to the physiological response time (*τ*). Consequently, protocols with similar nominal ramp rates but different dwell durations can produce fundamentally different physiological responses. Conceptually, this exposure structure is analogous to sequential transitions between fixed environments, rather than continuous exposure within a single gradually changing environment. **Figure 1** provides a schematic overview of both protocols and the corresponding wet-bulb temperatures.

**Figure 1.**
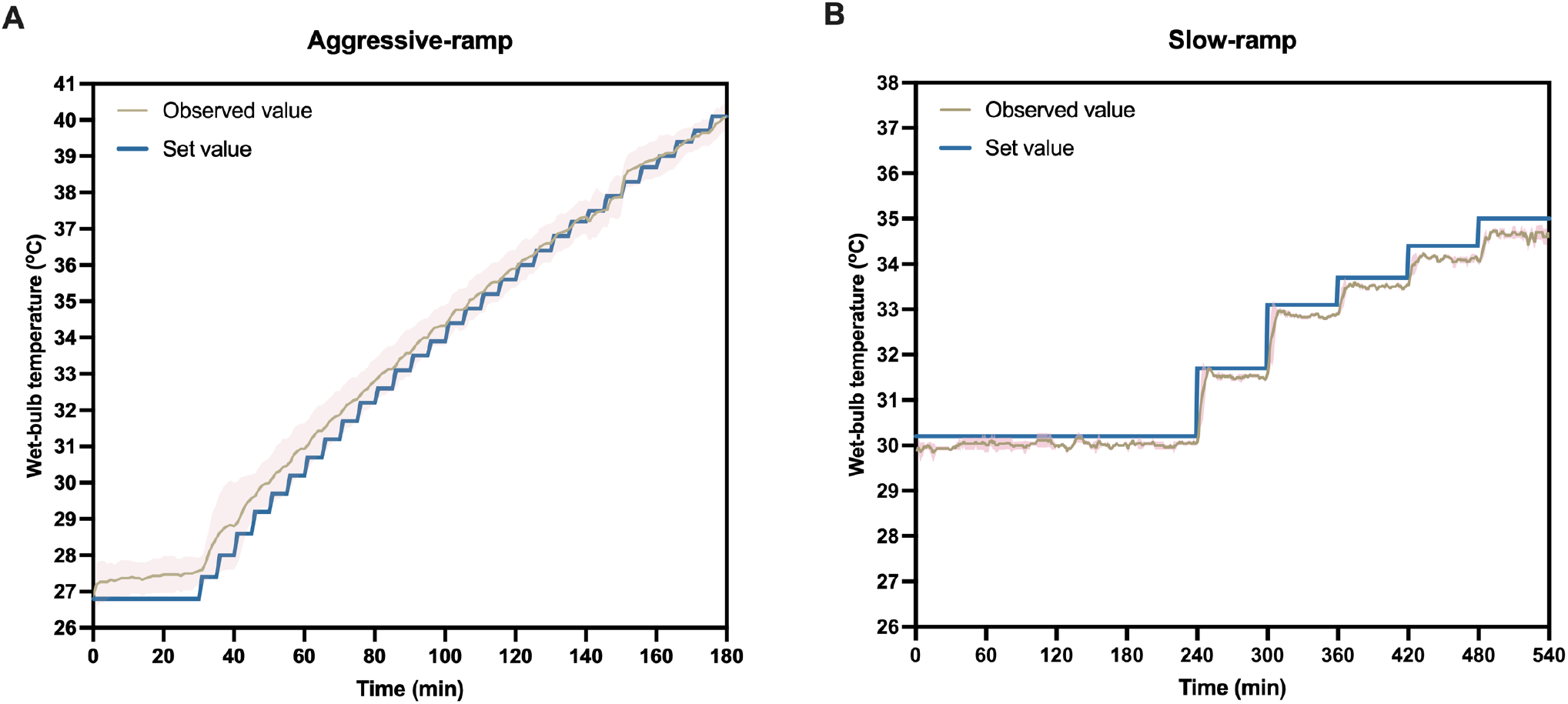
Schematic illustration of the two humidity ramp protocols. **A**, *Aggressive-ramp*: an initial 30-min stabilization at 42 °C, followed by 2% RH increments every 5 min until 88% RH (total exposure duration = 180 min); **B**, *Slow-ramp*: an initial 4-hour stabilization at 42 °C and 40% RH, followed by 6% RH increments per hour for 2 hours (to 52% RH), and then 3% RH increments per hour for up to 3 hours (to 61% RH; total exposure duration = 9 hours). The programmed set values (blue lines) and the observed mean wet-bulb temperature (*T*_w_) with standard deviation (tan line with pink band) are shown for each protocol. Note: all *T*_w_ values reported in the present study are calculated using the Stull equation (28).

#### 2.2.4 Measurements

Hydration status was assessed using a digital hand-held refractometer (PAL-1, Atago Co. Ltd., Tokyo, Japan; accuracy: ±0.2%). Rectal temperature (*T*_*cr*_) was measured using a thermistor probe (YSI-401, YSI Inc., Yellow Springs, OH) inserted 10 cm beyond the anal sphincter. Mean skin temperature (*T*_*sk*_) was obtained using iButtons (DS1992L, Maxim Integrated, Sunnyvale, CA; calibrated to accuracy: ±0.2 °C) affixed at the chest, upper arm, thigh, and calf, weighted according to Ramanathan’s four-site equation (29). Heart rate (HR) was continuously recorded using chest-strap telemetry (Polar Vantage XL, Polar Electro Oy, Finland). Metabolic rate was assessed using a breath-by-breath metabolic system (Calibre Biometrics, Wellesley, MA) for 10 minutes after a 15-minute thermal stabilization period during which participants performed light office tasks in the heat chamber (30). Perceptual ratings of thermal sensation, thermal discomfort, skin wetness, and thirst sensation were obtained every 5–10 min using validated scales (**Figure S1**) (31). Environmental conditions (*T*_db_ and RH) were continuously monitored using three calibrated temperature–humidity data loggers (TH42W-EX, HHW Technology Co. Ltd., Shenzhen, China; temperature accuracy: ±0.1 °C; humidity accuracy: ±1.5%). The loggers were positioned at heights of 0.1 m, 0.6 m, and 1.1 m above the floor. All physiological variables were recorded at 1-minute intervals.

#### 2.2.5 Termination criteria

Trials were immediately terminated if any of the following occurred: 1) *T*_*cr*_ reached or exceeded 38.6 °C; 2) onset of heat-related symptoms, including but not limited to involuntary hyperventilation, sustained piloerection, heat-related tetany, numbness, dizziness, nausea, confusion, or presyncope, as identified through participant self-report or clinical observation by a certified nurse; 3) voluntary withdrawal by the participant; or 4) the total exposure duration reached 9 hours for slow-ramp and adaptive-ramp conditions, or 3 hours for aggressive-ramp conditions, reflecting the intensity and rate of humidity increase in each protocol. Following termination, participants were immediately removed to a cool environment (21–24 °C) for monitored recovery.

#### 2.2.6 Data analysis

Individual *T*_*cr*_ data were smoothed using the *smooth* function with a smoothing parameter (*spar*) of 0.9 (17). The time derivative of *T*_*cr*_ (°C/h) was computed to identify inflection points. The critical environmental limit (CEL) was defined as the environmental condition corresponding to the onset of a sustained increase in *T*_*cr*_. Differences in the apparent critical wet-bulb temperature (*T*_w,critical_) between aggressive- and slow-ramp protocols were assessed separately in females (*n* = 12) and males (*n* = 14) using paired-samples t-tests. Sex differences in *T*_w,critical_ and body surface area-to-mass ratio were examined separately for both protocols using independent-samples *t*-tests with Welch’s correction. Effect sizes were calculated as Cohen’s *d*. All tests were two-tailed with *α* = 0.05 and all statistical analyses were conducted in IBM SPSS Statistics (version 29.0.1.0), with significance set at *P* < 0.05. Data are presented as mean ± SD (standard deviation) unless otherwise specified.

#### 2.2.7 Inflection point determination

Previous studies (8–16) identified the compensable–uncompensable transition visually based on trace curvature in *T*_*cr*_. Here, the inflection point was defined objectively as the first time the smoothed *T*_*cr*_ slope ≥0.25 °C/h and remained elevated thereafter. This criterion improves reproducibility and identifies the onset of sustained heat strain as reflected by a continuous increase in *T*_*cr*_. To assess robustness, sensitivity analyses were conducted using alternative slope thresholds (0.15, 0.20, 0.25, and 0.30 °C/h; see **Table S1** in the Supporting Information). Results confirmed that 0.25 °C/h provided stable and physiologically consistent estimates, whereas looser or stricter thresholds either advanced the transition unrealistically or delayed it toward higher core temperatures, potentially obscuring the early onset of uncompensable heat stress.

#### 2.2.8 Validation of CELs derived from slow- and adaptive-ramp protocols

To validate CEL estimates, prolonged fixed-condition exposures were conducted at *T*_db_=42 °C using independent healthy cohorts. Fourteen males (age: 24.4±1.8 yr; height: 1.76±0.04 m; body mass: 70.3±4.9 kg; BMI: 22.8±1.5 kg/m^2^) and fourteen females (age: 23.0 ± 1.8 yr; height: 1.59 ± 0.06 m; body mass: 51.3 ± 5.7 kg; BMI: 20.4 ± 2.2 kg/m^2^) were recruited under the same inclusion criteria and ethical approvals as those applied in the main study.

For males, exposures targeted *T*_w_ = 32.7 °C (expected compensable) and *T*_w_ = 34.3 °C (expected uncompensable); for females, target *T*_w_ values were 33.1 °C (expected compensable) and 34.6 °C (expected uncompensable). Participants remained seated in light clothing with *ad libitum* electrolyte intake, and trials lasted up to 8 hours or until predefined safety endpoints were reached (*T*_cr_ ≥ 38.6 °C, heart rate ≥ 90% age-predicted maximum, severe discomfort or heat-related tetany, or voluntary withdrawal). Rectal temperature (*T*_cr_), skin temperature, heart rate, and perceptual scales were monitored continuously.

Compensability was defined as stabilization of *T*_cr_ (slope < 0.10 °C/h after 3 hours, sustained for ≥ 20 min), whereas uncompensability was defined as a sustained rise in *T*_cr_ (slope > 0.10 °C/h) without evidence of plateau. These slope-based criteria are consistent with prior work and have been shown to be robust across threshold ranges (17, 22, 31).

## 3 Results

### 3.1 Physiological interpretation of the model

The model conceptualizes the human thermoregulatory system as a first-order dynamic process governed by a single thermal time constant (*τ*), which defines the rate at which the body approaches thermal equilibrium. Two contrasting response regimes emerge depending on the relationship between the dwell interval (Δ*t*) and this physiological time constant.

When the dwell interval is much shorter than the body’s thermal response time (Δ*t* ≪*τ*), core temperature does not approach its step-specific equilibrium before the next increment occurs. As a result, deviations between the environment-defined equilibrium temperature and the observed *T*_*cr*_ persist and accumulate across successive steps. This leads to a progressive increase in the rate of *T*_*cr*_ rise, producing a premature, apparent inflection point that may be misinterpreted as a physiological limit of heat tolerance. This acceleration reflects cumulative non-equilibrium dynamics arising from incomplete equilibration (see Discussion for mechanistic interpretation). Conversely, when the dwell interval is sufficiently long relative to the time constant (Δ*t* ≳*τ*), *T*_*cr*_ approaches near-equilibrium conditions within each step, thereby minimizing transient deviations. Under these conditions, the temporal evolution of core temperature more closely follows the equilibrium trajectory, and inflection points, if present, more accurately reflect the transition to uncompensable heat stress.

### 3.2 Empirical derivation of CELs from aggressive- and slow-ramp protocols

Critical environmental limits (CELs), expressed as critical wet-bulb temperature (*T*_w,critical_), differed substantially as a function of dwell duration (Δ*t*). Core temperature increased more rapidly and reached critical levels earlier during the aggressive-ramp protocol than during the slow-ramp protocol.

Across the 14 male participants, mean *T*_w,critical_ in the aggressive-ramp condition (Δ*t* = 5 min) was 29.9±1.6 °C, whereas the slow-ramp condition (Δ*t* = 60 min) yielded a significantly higher value of 33.4±0.5 °C (mean difference: 3.4±1.9 °C; *P* < 0.001, Cohen’s *d* = 1.78; **Figures 2A–2B** and **S2**–**S3**). Similarly, in females (*n* = 12), *T*_w,critical_ was significantly lower in the aggressive-ramp (30.3±0.9 °C) than in the slow-ramp condition (33.8±0.5 °C, mean difference: 3.5±0.9 °C; *P* < 0.001; Cohen’s *d* = 3.84; **Figures 2C–2D** and **S2**–**S3**). These findings demonstrate a strong dependence of apparent CELs on ramp dwell duration, with shorter intervals yielding systematically lower *T*_w,critical_ values. Consistent with model predictions, short dwell durations (Δ*t*/*τ* ≪1) were associated with a progressive increase in the rate of *T*_*cr*_ rise and earlier detection of inflection points. In contrast, longer dwell intervals allowed *T*_*cr*_ to approach near-equilibrium conditions within each step, resulting in higher and more stable *T*_w,critical_ estimates.

**Figure 2.**
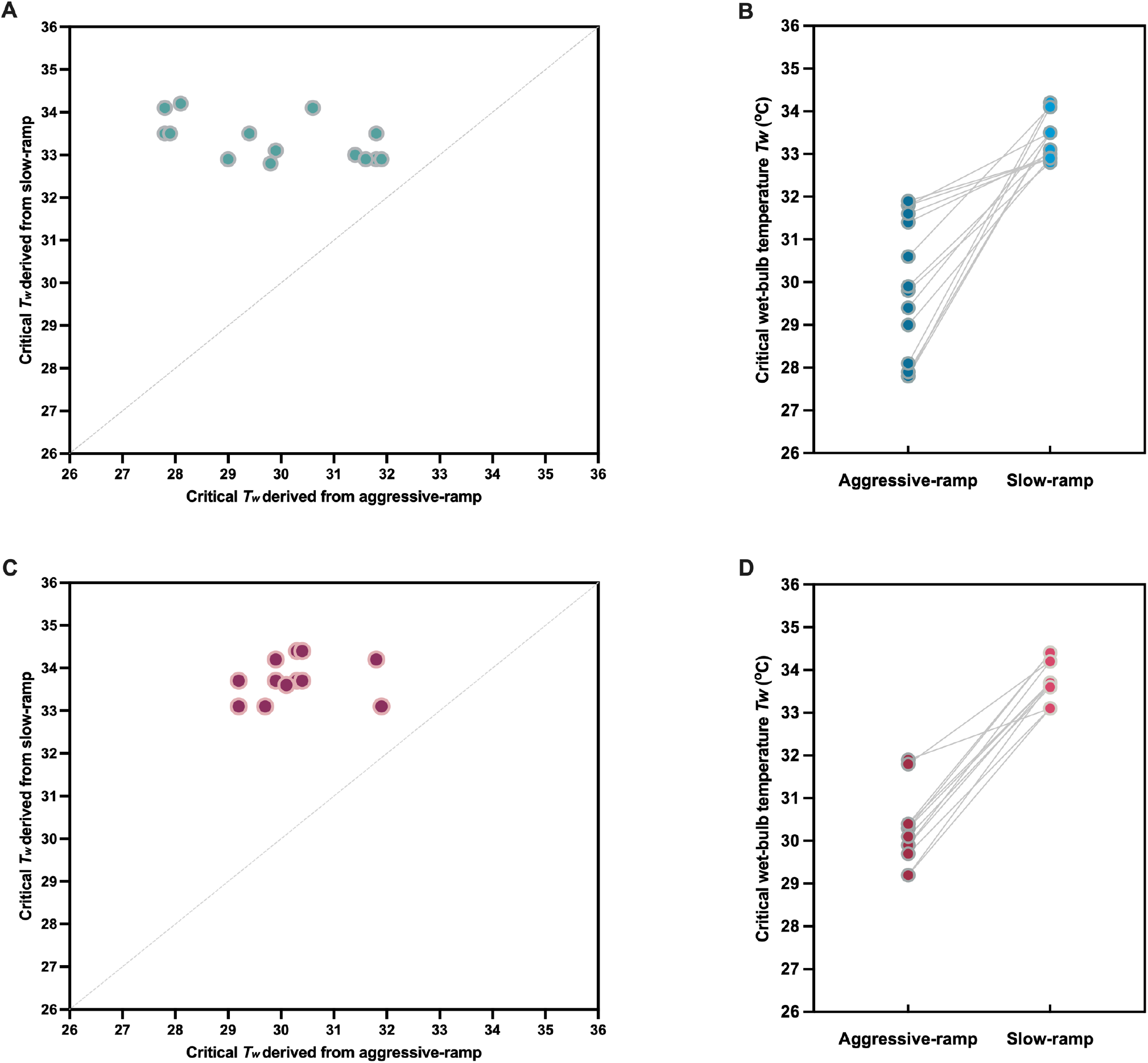
Critical wet-bulb temperatures (*T*_*w,critical*_) thresholds determined using aggressive- and slow-ramp humidity protocols at a constant dry-bulb temperature of 42 °C in males (*n*=14; **Panels A** and **B**) and females (*n*=12; Panels **C** and **D**). Panels **A** and **C** show scatter plots comparing *T*_w,critical_ values obtained from the aggressive-ramp (Δ*t* = 5 min) and slow-ramp (Δ*t* = 60 min) protocols for each participant. The grey dotted line represents the line of identity (*y* = *x*). All data points lie above this line, indicating systematically lower *T*_w,critical_ values with the aggressive-ramp protocol in both sexes. These systematic underestimations likely arise from the shorter ramp duration (2% RH increase per 5 min), which induces a premature inflection point compared with the slower protocol (60 min per 6%/3% increase). Panels **B** and **D** display paired matchstick plots, with each line connecting the same participant’s *T*_w,critical_ values from both conditions. All lines slope upward, confirming higher critical limits with the slow-ramp protocol. The narrower distribution of slow-ramp values indicates lower inter-individual variability and greater measurement consistency, whereas the wider spread in aggressive-ramp values reflects higher inter-individual variability under rapid humidity increments in both sexes.

Under the aggressive-ramp protocol, no significant sex difference in *T*_w,critical_ was observed (females: 30.3±0.9 °C; males: 29.9±1.6 °C; Welch’s *t*(20.2) = 0.69, two-tailed *P* = 0.497, one-tailed *P* = 0.248, Cohen’s *d* = 0.26). In contrast, under the slow-ramp protocol, females exhibited higher *T*_w,critical_ values than males (33.8±0.5 °C vs. 33.4±0.5 °C), with a large effect size (Cohen’s *d* = 0.80). This difference approached significance in a two-tailed test (Welch’s *t*(23.6) = 2.03, *P* = 0.053) and reached significance in a one-tailed test (*P* = 0.027), consistent with the a-priori directional hypothesis.

To further characterize equilibration dynamics, we implemented an **adaptive-ramp protocol** in which relative humidity (RH) increments (3%, 6% or 10%) were applied only after full initial equilibration of *T*_*cr*_. **Figure 3** illustrates representative rectal temperature (*T*_*cr*_) responses from six male and three female participants at *T*_db_=42 °C. Across participants, re-establishment of near steady-state *T*_*cr*_ following RH increments required substantially longer durations than the nominal environmental transition time. After 3% RH increments, equilibration occurred approximately 38–91 min in some individuals, but often required over 200–400 min following successive steps. Larger increments (6– 10% RH) further prolonged equilibration, with stabilization times ranging from ~98 to >200 min. In several cases (e.g., **Figures 3A** and **3E**), no inflection point was observed even after prolonged exposure (>9 h), indicating sustained compensability. These observations demonstrate that, although chamber humidity reached target levels within 2–5 min, physiological equilibration of *T*_*cr*_ occurred on timescales an order of magnitude longer. Consequently, conventional ramp protocols with 5–10 min dwell durations (Δ*t*/*τ* ≪1) are insufficient to permit equilibration and therefore systematically underestimate true CELs.

**Figure 3.**
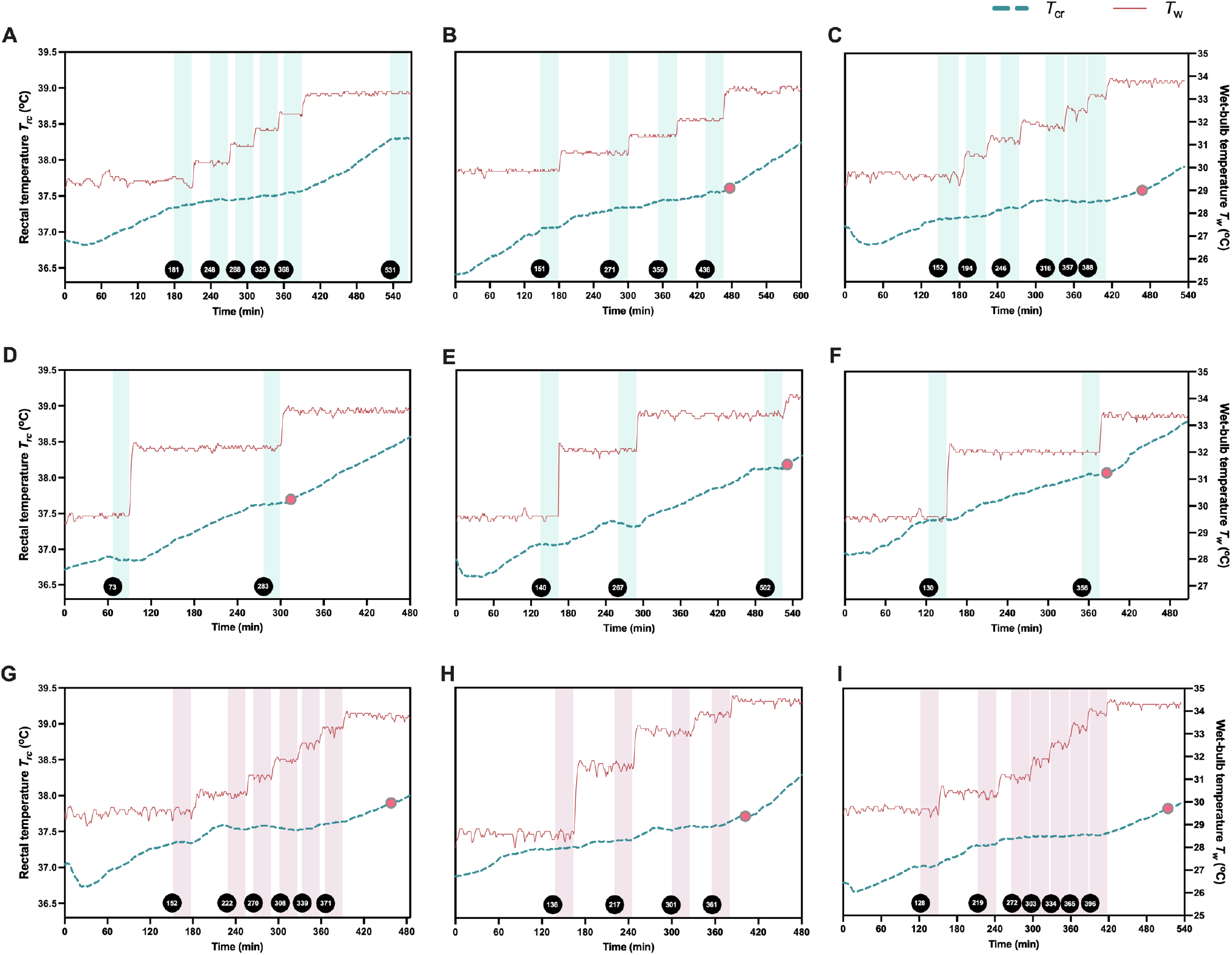
Time course of rectal temperature (*T*_*cr*_) re-stabilization under the adaptive ramp protocol in six representative male (**A**–**F**) and three female (**G**–**I)** participants at *T*_db_=42 °C, following initial equilibration at 38% RH. Stepwise relative humidity (RH) increments of 3%, 6%, or 10% were applied only after prior *T*_*cr*_stabilization (defined as |d*T*_*cr*_ /d*t*| ≤ 0.1 °C over 30 min). Green and pink shaded areas denote the 30-min stabilization periods in males and females, respectively. Total exposure durations ranged from 8–10 h (480–600 min). Circles with a grey border and pink fill mark the identified inflection points. No inflection was observed in panel **A** throughout the entire 570 min. Inflection points in panels **B**–**I** occurred at 33.4 °C (447 min), 33.7 °C (476 min), 33.5 °C (319 min), 34.0 °C (536 min), 33.5 °C (392 min), 33.8 °C (465 min), 34.5 °C (409 min), and 34.3 °C (520 min), respectively.

Inflection points identified under the adaptive ramp protocol (males: 33.4–34.0 °C; females: 33.8– 34.5 °C; **Figure 3**) closely matched those obtained under the slow ramp protocol (males: 33.4±0.5 °C; females: 33.8±0.5 °C) and were substantially higher than values derived from the aggressive-ramp protocol. Although full stabilization was not always achieved in the slow-ramp protocol, the resulting *T*_w,critical_ values remained consistent with those obtained under adaptive-ramp conditions. Collectively, these findings indicate that slow-ramp protocols provide substantially more accurate estimates of physiological heat tolerance, consistent with those observed under prolonged fixed-condition exposures (21, 22, 31).

### 3.3 CEL validation against prolonged fixed-condition exposures

Prolonged fixed-condition exposure trials at *T*_db_ = 42 °C confirmed the validity of CELs derived from the slow- and adaptive-ramp protocols. In males (*n* = 14), exposures at *T*_w_ = 32.6±0.4 °C resulted in stabilization of *T*_cr_ (slope < 0.10 °C/h after 5.8 hours), indicating compensable heat stress, whereas *T*_w_ = 34.3±0.2 °C produced a sustained increase in *T*_cr_ (slope > 0.10 °C/h without plateau), consistent with uncompensable conditions (**Figures 4A–4B**). Similarly, in females (*n* = 14), *T*_w_ = 32.9±0.4 °C was compensable, with near-stable *T*_cr_ dynamics (slope < 0.10 °C/h after 4.2 hours), whereas *T*_w_ = 34.6±0.4 °C resulted in progressive heat storage (**Figures 4C–4D**). These thresholds closely align with values derived from the slow-ramp (males: 33.4 °C [32.8–34.2 °C]; females: 33.8 °C [33.0–34.3 °C]) and adaptive-ramp protocols (males: 33.4–34.0 °C; females: 33.8–34.5 °C), supporting their physiological relevance. In contrast, the lower *T*_w,critical_ values obtained under the aggressive-ramp protocols (males: ~29.9 °C; females: ~30.3 °C) fall well below the compensable range identified under prolonged exposure conditions, indicating a systematic underestimation of CELs.

**Figure 4.**
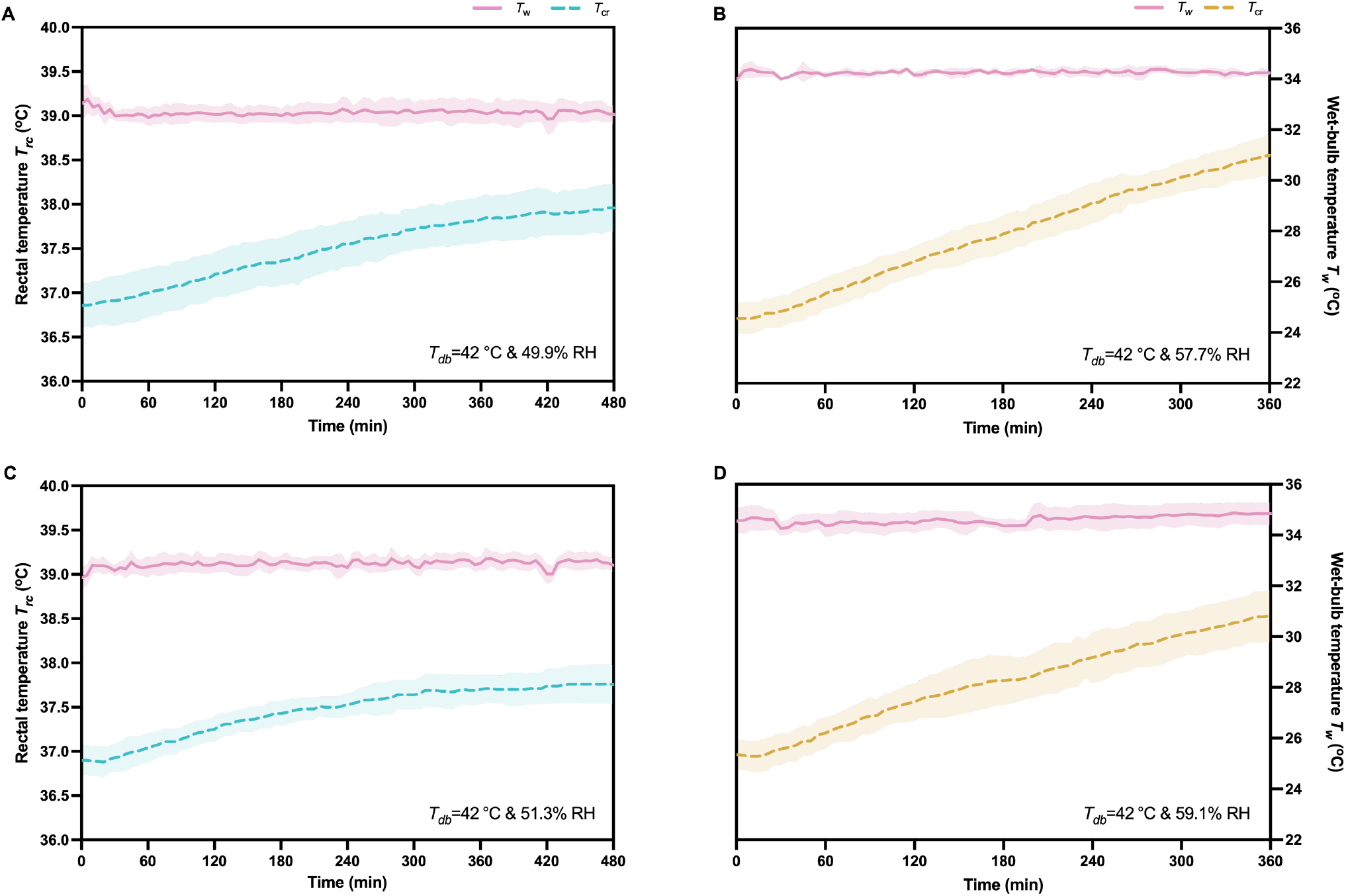
Time course of rectal temperature (*T*_*cr*_) responses and ambient wet-bulb temperature (*T*_w_) during prolonged fixed-condition exposures at *T*_db_=42 °C in males and females. A, *T*_*cr*_ in males at target *T*_w_=32.7 °C (compensable) and *T*_w_=34.3 °C (uncompensable); B, corresponding measured *T*_w_ in the heat chamber for males; C, *T*_*cr*_ in females at target *T*_w_=33.1 °C (compensable) and *T*_w_=34.6 °C (uncompensable); D, corresponding measured *T*_w_ in the heat chamber for males; Data are means ± SD; compensability defined as *T*_*cr*_ stabilization (< 0.10 °C/h slope after 3 h, maintained ≥20 min), uncompensability as sustained *T*_*cr*_ rise (> 0.10 °C/h without plateau).

### 3.4 Sensitivity analysis

The first-order dynamical model indicates that the ratio Δ*t*/*τ* (i.e., step dwell time relative to the physiological time constant) governs whether *T*_*cr*_ can approach equilibrium at each stage. When Δ*t*/*τ* ≪1, equilibration is incomplete and residual deviations from equilibrium persist across successive steps, resulting in a progressive increase in the rate of *T*_*cr*_ rise and earlier identification of inflection points.

Empirically derived estimates of the core temperature time constant (*τ*), obtained following initial equilibration, ranged from approximately 1.0 to >3.3 hours, depending on metabolic rate, hydration status, and thermoregulatory efficiency. Under these conditions, step durations of 5–10 min, commonly used in ramp-based protocols, are substantially shorter than the physiological response timescale (8, 9, 11, 13, 16). Accordingly, such protocols operate in a regime where Δ*t*/*τ* ≪1, and near-equilibrium conditions are not achieved within each step, biasing the apparent location of the inflection point. We therefore recommend either empirical estimating *τ* for the study population or performing sensitivity analyses across plausible *τ* values to ensure robustness of CEL estimates. **Table 1** summarizes the dependence of thermoregulatory system behavior on Δ*t*/*τ*.

**Table 1.** Sensitivity analysis of the dynamic response to step duration (*Δt*) relative to the physiological time constant (*τ*). CELs, critical environmental limits.

| $\Delta t$<br>(min) | Physiological time constant ($\tau$, min) | | | Predictive response characteristic | Impact on CELs |
| --- | --- | --- | --- | --- | --- |
|  | 120 | 180 | 240 |  |  |
| 5 | 0.04 | 0.03 | 0.021 | High kinetic mismatch: environment changes much faster than heat loss can activate; pronounced false early $T_{cr}$ inflection | Underestimate |
| 10 | 0.08 | 0.06 | 0.042 | Moderate-to-high kinetic mismatch: environment changes markedly faster than heat loss activation; notable false early $T_{cr}$ inflection | Underestimate |
| 60 | 0.5 | 0.33 | 0.25 | Low kinetic mismatch: environment changes at a rate comparable to heat loss activation; partial equilibration with minimal false inflection | Near true value |
| 120 | 1.0 | 0.67 | 0.50 | Very low kinetic mismatch: environment changes slower than or equal to heat loss activation; full equilibration, no false inflection | Accurate |

**Note:** When the ramp step duration ( $\Delta t$ ) is much shorter than the physiological time constant ( $\tau$ ), the rate of environmental change outpaces the finite activation kinetics of heat loss responses. This kinetic mismatch prevents the thermoregulation system from reaching its required compensatory capacity, leading to a false early inflection in core temperature and a systematic underestimation of the critical environmental limit (CEL). The key determinant is the dimensionless ratio $\Delta t/\tau$ . When $\Delta t/\tau \ll 1$ , the environment outruns thermoregulatory kinetics, causing a false early inflection that leads to systematic underestimation of the true CEL. When $\Delta t/\tau \gtrsim 0.3\text{--}0.5$ , near steady-state conditions are achieved, yielding accurate CEL estimates.

## 4 Discussion

This study demonstrates that apparent critical environmental limits (CELs; i.e., *T*_w,critical_values) derived from humidity-ramp protocols are strongly dependent on the temporal structure of environmental change, rather than reflecting fixed physiological thresholds. Specifically, short dwell durations systematically yield lower *T*_w,critical_ values, whereas slow- and adaptive-ramp protocols produce estimates that closely align with those obtained from prolonged fixed-condition exposures. These findings indicate that commonly used ramp protocols introduce a systematic downward bias in inferred heat tolerance limits.

A key conceptual distinction underlying the present work is that humidity-ramp protocols do not measure the steady-state relationship between environment and core temperature, but rather the transient trajectory of a dynamic system under continuously shifting boundary conditions. In the absence of thermal lag (i.e., *τ* = 0, [32, 33]), core temperature would instantaneously track the equilibrium value associated with each environmental step, and the timing of step transitions (Δ*t*) would be irrelevant. Under such conditions, no kinetic mismatch would arise, and all ramp protocols would converge to the same critical environmental limit (CEL). However, in biological systems, thermal lag (τ > 0), determined by tissue heat capacity, heat exchange processes, and physiological regulation (32–37) introduces a finite response time, such that core temperature evolves toward — rather than instantaneously reaches — the equilibrium state. When environmental conditions change more rapidly than this response timescale (17–20, 31, 38), the system remains in a non-equilibrium state, such that *T*_cr_ continuously adjusts toward a shifting equilibrium rather than stabilizing at each step. Importantly, although thermal lag delays the approach to equilibrium under fixed conditions, in ramp protocols it advances the apparent inflection point because the system is evaluated against a continuously shifting equilibrium that it never reaches. More specifically, the observed inflection in *T*_cr_ does not indicate that the current environment has exceeded the body’s true heat dissipation capacity. Instead, it reflects a dynamic mismatch between the rate of environmental forcing and the kinetics of physiological adjustment. As a result, the inflection point occurs before the system has had sufficient time to approach the equilibrium state corresponding to that environment, leading to a systematic downward bias in apparent CEL estimates.

Consistent with this framework, our results show that when dwell durations are much shorter relative to the physiological response time (Δ*t*/*τ* ≪1), deviations from equilibrium persist and accumulate across successive steps, producing a progressive increase in the rate of *T*_cr_ rise and premature identification of inflection points. In contrast, when dwell durations are sufficiently long (Δ*t*/*τ* ≳0.3–0.5), *T*_cr_ approaches near-equilibrium conditions within each step, yielding smoother trajectories and more stable estimates of *T*_w,critical_. This dimensionless ratio (Δ*t*/*τ*) therefore provides a generalizable criterion for evaluating ramp adequacy across experimental designs and populations.

A key methodological implication is that humidity-ramp protocols, although often described as continuous environmental changes, are implemented in practice as discrete step transitions. Each step imposes a new equilibrium condition, and the physiological response is governed primarily by the dwell duration at that condition. When this duration is insufficient relative to *τ*, incomplete equilibration is unavoidable, and the resulting *T*_cr_ trajectory reflects cumulative non-equilibrium dynamics. This provides a mechanistic explanation for why ramp-derived inflection points may occur at environmental conditions that remain physiologically compensable under prolonged exposure.

This framework also resolves longstanding discrepancies in the literature. Belding and Kamon (8) recognized that rectal temperature may require “*more than 2 hours to re-stabilize*” under critical conditions, yet subsequent shortcut ramp protocols increased humidity at fixed short intervals following only an initial equilibration period. While this approach improved feasibility, it did not account for the finite response time required after each environmental increment. Our empirical data show that even modest humidity increases (e.g., 3%) require ~0.6–1.5 h for *T*_*cr*_ re-stabilization (**Figure 3**), far exceeding the 5–10 min dwell durations commonly used. Consequently, previously reported *T*_w,critical_ values of 26.0–32.3 °C likely reflect kinetically constrained, non-equilibrium dynamics rather than true physiological limits (10, 11, 13–16). This mechanism reconciles these lower estimates with steady-state-derived thresholds of ~33–34 °C (22, 31, 39), with the ~3–4 °C discrepancy arising from cumulative incomplete equilibration at *T*_db_ = 42 °C.

Further support is provided by the adaptive ramp protocol. When each humidity increment was maintained until *T*_*cr*_ approached a near steady state, the resulting *T*_w,critical_ values (males: 33.4–34.0 °C; females: 33.8–34.5 °C) closely matched those obtained from both the slow-ramp protocol and prolonged fixed-condition exposures. In contrast, aggressive-ramp protocols systematically underestimated these thresholds and misclassified compensable environments as uncompensable. At *T*_db_ = 42 °C, each 3% RH increment corresponds to ~0.7 °C increase in *T*_w_, such that rapid step progression can traverse the narrow transition zone between compensable and uncompensable heat stress (~33–34 °C) without allowing sufficient time for equilibration.

Comparison between aggressive (Δ*t* = 5 min) and slow (Δ*t* = 60 min) ramp protocols further underscores the importance of temporal pacing. Slow-ramp-derived *T*_w,critical_ values (33.4 °C in males and 33.8 °C in females) were independently validated using prolonged fixed-condition exposures and align closely with previously reported steady-state thresholds in young adults (22, 31, 39). In contrast, aggressive-ramp protocols operate in a regime where Δ*t*/*τ* ≪1, such that *T*_cr_ reflects a transient trajectory rather than a steady-state response. This leads to earlier inflection points and systematically lower apparent CELs.

Interestingly, no significant sex difference in *T*_w,critical_ was observed under the aggressive-ramp protocol, likely because rapid environmental change maintains the system in a strongly non-equilibrium state in which transient heat storage dominates. Under the slow-ramp protocol, however, females exhibited a modest but consistent advantage (~0.4–0.5 °C advantage). This difference is attributable to morphological and biophysical factors, including a higher surface area-to-mass ratio (0.0287±0.0018 m^2^/kg [females] vs. 0.0257±0.0012 m^2^/kg [males], two-sided *P*<0.001; Cohen’s *d* = 2.023) and lower absolute body mass (females: 55.3±6.1 kg; males: 74.9±9.2 kg), which enhance relative evaporative heat loss under humid conditions. These findings are consistent with prior work indicating that sex differences in heat tolerance become more apparent when steady-state heat balance is approached (12, 13, 40).

Collectively, these results indicate that apparent sex differences in heat tolerance are strongly time-scale dependent and emerge only under conditions that permit near-equilibrium thermoregulation. Females generally exhibit lower absolute sweat rates but greater evaporative efficiency per unit surface area, resulting in reduced “wasted” sweat under high humidity conditions. Such biophysical advantages have long been proposed to favor females during prolonged warm-humid exposures when fitness and acclimatization are comparable (7, 40). More broadly, they highlight that experimental design—not just physiology—can shape inferred heat tolerance limits. Future work across diverse populations, activity levels, and environmental conditions will be essential to disentangle the relative contributions of morphology, physiology, and protocol design.

### 4.1 Physiological and methodological implications

We identify the ratio Δt/τ as a dimensionless similarity criterion governing heat tolerance assessment, where Δ*t* represents the dwell duration of each humidity step and *τ* is the physiological time constant describing the rate at which rectal temperature (*T*_cr_) approaches its environment-defined equilibrium. This ratio provides a quantitative framework for determining whether thermophysiological responses can adequately track imposed environmental changes. When Δ*t*/*τ* < 0.2, the system remains far from equilibrium and ramp protocols induce substantial dynamic artifacts due to incomplete adjustment. Values of 0.2 ≤ Δ*t*/*τ* < 0.5 permit partial equilibration, whereas Δ*t*/*τ* ≥ 0.5 approaches quasi–steady-state conditions with minimal distortion of the *T*_cr_ trajectory. Empirically, a ratio of ~0.3 was sufficient to achieve partially stable core temperature responses without excessively prolonging exposure. Estimating *τ* from the exponential rise of *T*_*cr*_ during the initial equilibration phase enables individualized ramp pacing aligned with thermophysiological response kinetics.

This framework explains why rapid humidity-ramp protocols consistently yield artificially low CELs and underscores the need for standardized ramp parameters across laboratories. When Δ*t*/*τ* ≪1, successive environmental steps occur before the system approaches the equilibrium associated with the preceding condition, leading to cumulative kinetically constrained, non-equilibrium dynamics. As a result, *T*_*c*_ trajectories become progressively steeper and inflection points emerge prematurely. Importantly, although such dynamics would delay equilibration under fixed environmental conditions, in ramp protocols they instead advance the apparent inflection point because the system is evaluated against a continuously shifting equilibrium that it does not reach. These inflection-based thresholds should therefore be interpreted as manifestations of transient system dynamics rather than steady-state physiological limits. In this context, prolonged fixed-condition exposures remain the definitive benchmark for defining true environmental compensability and for calibrating predictive models, occupational exposure limits (OELs), and wearable heat-strain algorithms (41).

The methodological implications are substantial. Conventional aggressive humidity-ramp protocols implicitly assume that thermophysiological responses can equilibrate within each step; however, our data demonstrate that typical dwell durations of 5–10 minutes are markedly shorter than the physiological response timescale (17, 18). For example, at *T*_*w*_≈ 31.7 °C (*T*_*db*_ = 42 °C), *T*_*cr*_ required ≥ 98 min (1.63 h) to re-establish near steady-state conditions following a 10% RH increment (**Figures 3D–3F**). Under such conditions, the observed inflection reflects a trajectory of ongoing adjustment toward a shifting equilibrium, rather than a true transition to uncompensable heat stress. This mismatch produces persistent positive heat storage, yielding progressively steeper *T*_cr_ trajectories and systematic underestimation of heat tolerance limits.

Accordingly, aggressive humidity-ramp protocols should be regarded as rapid but approximate tools for screening environmental compensability. Accurate estimation of CELs requires that each environmental step be maintained long enough to permit at least partial equilibration, as implemented in the slow- and adaptive-ramp protocols. While the adaptive ramp protocol provides the most precise characterization, some individuals may require exposure durations exceeding 10 hours to reach an inflection point. The slow-ramp protocol therefore represents a practical compromise between physiological validity and experimental feasibility. Nevertheless, both approaches require validation against prolonged fixed-condition exposures, which minimize transient effects and capture true steady-state limits. Prolonged fixed-condition exposure thus remains the methodological benchmark for quantifying human heat tolerance, whereas ramp-based approaches — when appropriately designed — provide rapid but approximate estimates of environmental compensability, and should not be interpreted as direct measures of steady-state heat tolerance unless near-equilibrium conditions are achieved (17, 42, 43).

### 4.2 Limitations and future directions

This study identified a fundamental methodological artifact in conventional humidity-ramp protocols using a relatively homogeneous cohort of young, healthy adults at a fixed dry-bulb temperature. This controlled design was intentional, enabling isolation of the effects of ramp timing (Δ*t*) on thermophysiological responses while minimizing inter-individual variability. However, it necessarily limits the generalizability of the absolute critical environmental limit (CEL) values reported here, given the well-established influences of age, acclimatization status, fitness, and health on thermoregulatory capacity (43, 44).

Importantly, the artifact identified in this study, namely, the premature appearance of a *T*_*cr*_inflection point under short dwell durations, is not population-specific, but reflects a fundamental constraint imposed by the finite kinetics of human thermoregulation. The temporal evolution of *T*_*cr*_ is governed by a physiological time constant (*τ*), which limits the rate at which the body can approach equilibrium following environmental change (17). When environmental transitions occur on timescales shorter than this response time (Δ*t*/*τ* ≪1), the system necessarily operates under kinetically constrained, non-equilibrium conditions. Under such conditions, *T*_*cr*_ does not reflect the steady-state response to the current environment, but instead integrates residual effects of prior exposures. Consequently, the observed inflection point occurs before the system has approached the equilibrium corresponding to that environmental condition, leading to a systematic underestimation of the true CEL.

While absolute CEL values may differ across populations due to variations in morphology, evaporative capacity, cardiovascular function, and behavioral responses, the direction of bias introduced by insufficient dwell time is expected to be consistent. Specifically, short-dwell ramp protocols will systematically underestimate the true compensable–uncompensable boundary, because the underlying mechanism — a mismatch between the rate of environmental forcing and the kinetics of physiological adjustment — is universal.

The Δ*t*/*τ* framework proposed here provides a generalizable approach for addressing this limitation. By explicitly accounting for the relationship between environmental pacing and physiological response time, future studies can design ramp protocols that better approximate equilibrium conditions or apply post hoc corrections to existing datasets. This is particularly important for studies involving older adults, clinical populations, or other vulnerable groups, in whom longer response times (i.e., larger *τ*) may further amplify non-equilibrium effects. Incorporating this framework into experimental design and analysis will improve the accuracy, comparability, and physiological interpretability of human heat tolerance assessments across diverse populations.

## 5 Conclusions

Humidity-ramp protocols have long served as practical tools for assessing safe occupational exposure in hot environments and have recently been used to estimate critical environmental limits (CELs) of human heat tolerance. The present study demonstrates that these protocols systematically underestimate heat tolerance due to a mismatch between the rate of environmental forcing and the kinetic of physiological adjustment. When environmental changes occur on timescales shorter than the physiological response time (Δ*t*/*τ* ≪1), core temperature does not approach its step-specific equilibrium, and the resulting inflection points reflect transient, non-equilibrium dynamics rather than a true transition from compensable to uncompensable heat stress.

This kinetic constraint leads to a consistent downward bias in critical wet-bulb temperature estimates, on the order of ~3–4 °C compared with values derived from prolonged fixed-condition exposures at *T*_db_ = 42 °C in healthy young adults. Accordingly, aggressive-ramp protocols should not be used as standalone methods for defining absolute limits of human heat tolerance. Prolonged fixed-condition exposures remain the methodological benchmark for identifying true compensability boundaries. Where such approaches are impractical, slow-ramp protocols with sufficiently long dwell durations (e.g., ≥ 60 min) provide a more reliable alternative by allowing partial equilibration and reducing non-equilibrium artifacts. More broadly, the ratio Δ*t*/*τ* emerges as a governing dimensionless parameter for both interpreting existing data and designing future heat tolerance assessments. Explicit consideration of this relationship will improve the physiological validity of experimental protocols and enhance the accuracy of heat tolerance limits used to inform occupational standards and public health strategies under extreme heat.

## Notes

### Competing Interest Statement

The authors have declared no competing interest.

### Summary of Updates

The entire manuscript has been revised to more clearly articulate the mismatch between the rate of environmental forcing and the kinetics of physiological adjustment. The concept of thermal lag has been explicitly incorporated to explain why a dwell time of 5 &10 minutes is fundamentally insufficient for core temperature to stabilize during successive ramp steps. In addition, the manuscript has been substantially shortened throughout to improve clarity and conciseness.

### Funding

- National Excellent Young Scientist Program: 6119924022
- Sanqin Scholars: 2050225003

