## Supplemental figures and tables for "Revisiting the Humidity Ramp Protocol for Assessing Human Heat Tolerance Limits"

by

Fèlix Faming Wang, Yi Xu, Haojian Wang, Min Cui, Xue Hou, Boan Wei, and Xiong Shen

This file contains 7 figures and 2 tables.

**Mathematical justification**

The whole-body balance equation is read as

$$C\frac{dT_{cr}}{dt} = \dot{M} - \dot{W} + \dot{Q}_{res} - \dot{Q}_{skin}(T_{cr}) \quad \text{Eq.(S1)}$$

Define the net heat-loss function

$$\mathcal{L}(T_{cr}, t) \equiv \dot{Q}_{loss} = \dot{Q}_{conv}(T_{sk}, T_a) + \dot{Q}_{rad}(T_{sk}, T_r) + \dot{Q}_{evap}(p_s, p_a) + \dot{Q}_{res} \quad \text{Eq.(S2)}$$

where $T_{sk}$, $T_a$, $T_r$, $p_s$ and $p_a$ are themselves functions of $T_{cr}$ and physiological state.

At an operating point (instantaneous equilibrium) $T_{cr} = T_{cr,eq}$ then we have

$$\dot{M} - \dot{Q}_{loss}(T_{cr,eq}) = 0 \quad \text{Eq.(S3)}$$

For small perturbations $x(t) \equiv T_{cr}(t) - T_{cr,eq}$, perform a first-order Taylor expansion about $T_{cr,eq}$, yielding

$$\dot{M} - \dot{Q}_{loss}(T_{cr}) \approx -\frac{\partial \dot{Q}_{loss}}{\partial T_{cr}}\bigg|_{eq} x(t) + \mathcal{O}(x^2) \quad \text{Eq.(S4)}$$

where $\mathcal{O}(x^2)$ represents the error term due to truncating the Taylor expansion, which is proportional to $x^2$ and becomes negligible for small values of $x$. In our model, $\mathcal{O}(x^2)$ means that the dynamic lag introduced by linearizing $\dot{Q}_{loss}$ is small enough for small $\Delta T_{cr}$, and higher-order terms do not significantly affect the accuracy of core temperature predictions. The term $\frac{\partial \dot{Q}_{loss}}{\partial T_{cr}}\Big|_{eq} x(t)$ represents the rate change of heat loss $\dot{Q}_{loss}$ with respect to core temperature, evaluated at equilibrium. This term captures the sensitivity of heat loss to changes in body temperature. When the body's core temperature deviates from equilibrium (i.e., $x(t)=T_{cr}(t)-T_{cr,eq}$), this sensitivity determines the amount of heat dissipated by the body, driving the return to thermal equilibrium.

Substitute Eq.(S4) into whole-body balance Eq.(S1)

$$C\dot{x}(t) \approx -\frac{\partial \dot{Q}_{loss}}{\partial T_{cr}}\bigg|_{eq} x(t) \quad \text{Eq.(S5)}$$

This yields

$$C\dot{T}_{cr} = -k\left[T_{cr} - T_{cr,eq}\right] \quad \text{Eq.(S6)}$$

With the coefficient $k$ defined as

$$k \equiv -\frac{\partial \dot{Q}_{loss}}{\partial T_{cr}}\bigg|_{eq} \quad \text{Eq.(S7)}$$

The above linearization is valid when $|\ x\ |$ is small enough that second-order terms are negligible, i.e., when experiments probe dynamics near the instantaneous equilibrium.

**Perceptual responses**

Perceptual perceptions were registered at 5–10 min intervals. Rating scales included thermal sensation (9-point continuous scale: from -4 = very cold to +4 = very hot), thermal discomfort (7-point continuous scale, from -3 = very uncomfortable to +3 = very comfortable, thirst sensation (7-point continuous scale, from 1 =

not thirsty at all to 7 = very, very thirsty), and skin wetness sensation (7-point continuous scale, from -3 = very wet to +3 = very dry) (see **Figure S1**).

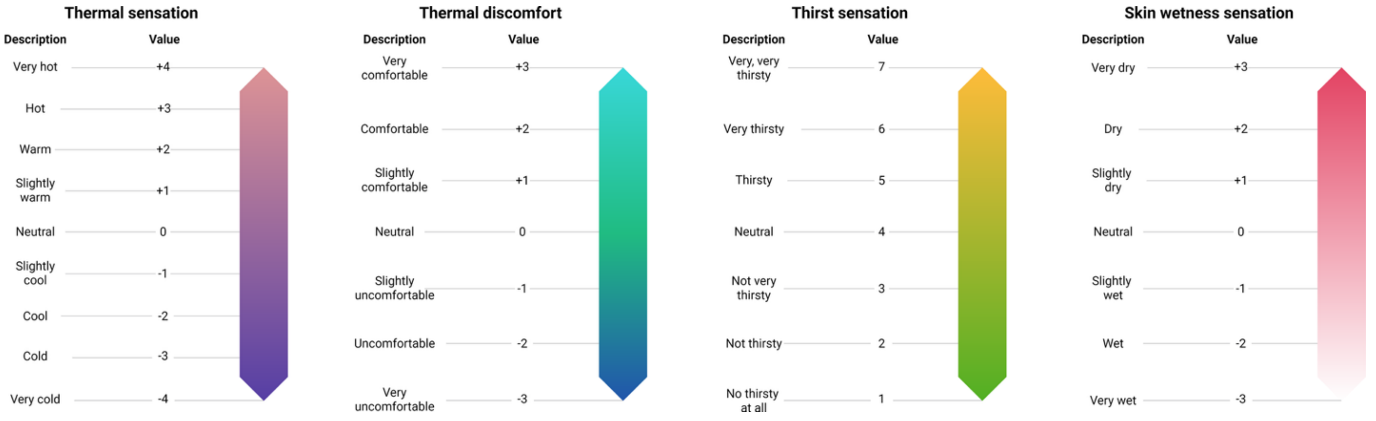


**Figure S1** Ratings scales for perceptual responses, including thermal sensation, thermal discomfort, thirst sensation, and skin wetness sensation.

### Supplemental figures

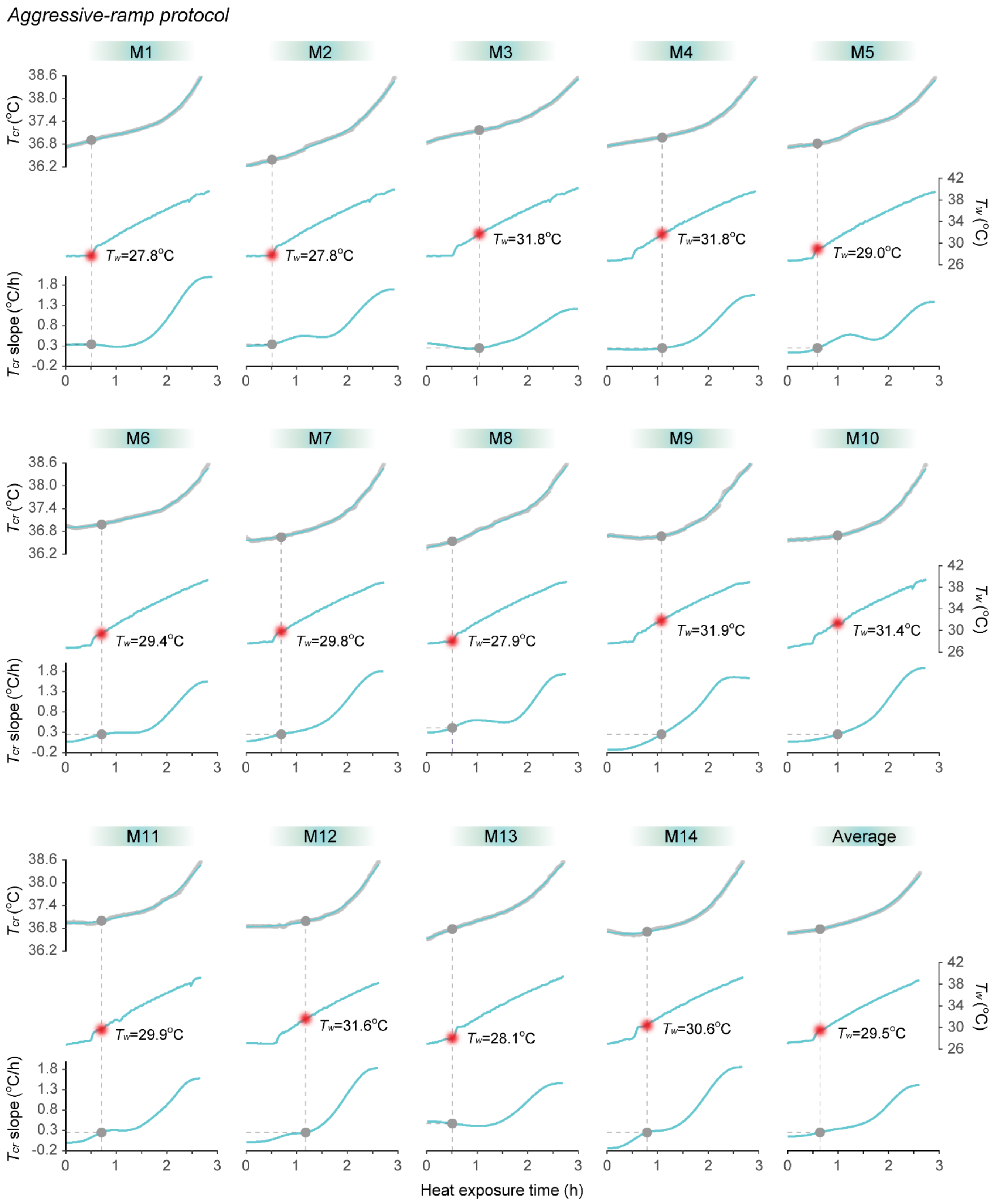


**Figure S2** Temporal variations in individual and average core temperature response, recorded environmental wet-bulb temperature in the heat chamber, and the slope of the rectal temperature changes under the aggressive-ramp protocol at a dry-bulb temperature of 42 °C in young adult males. M1–M14 denote male participants 1 through 14.

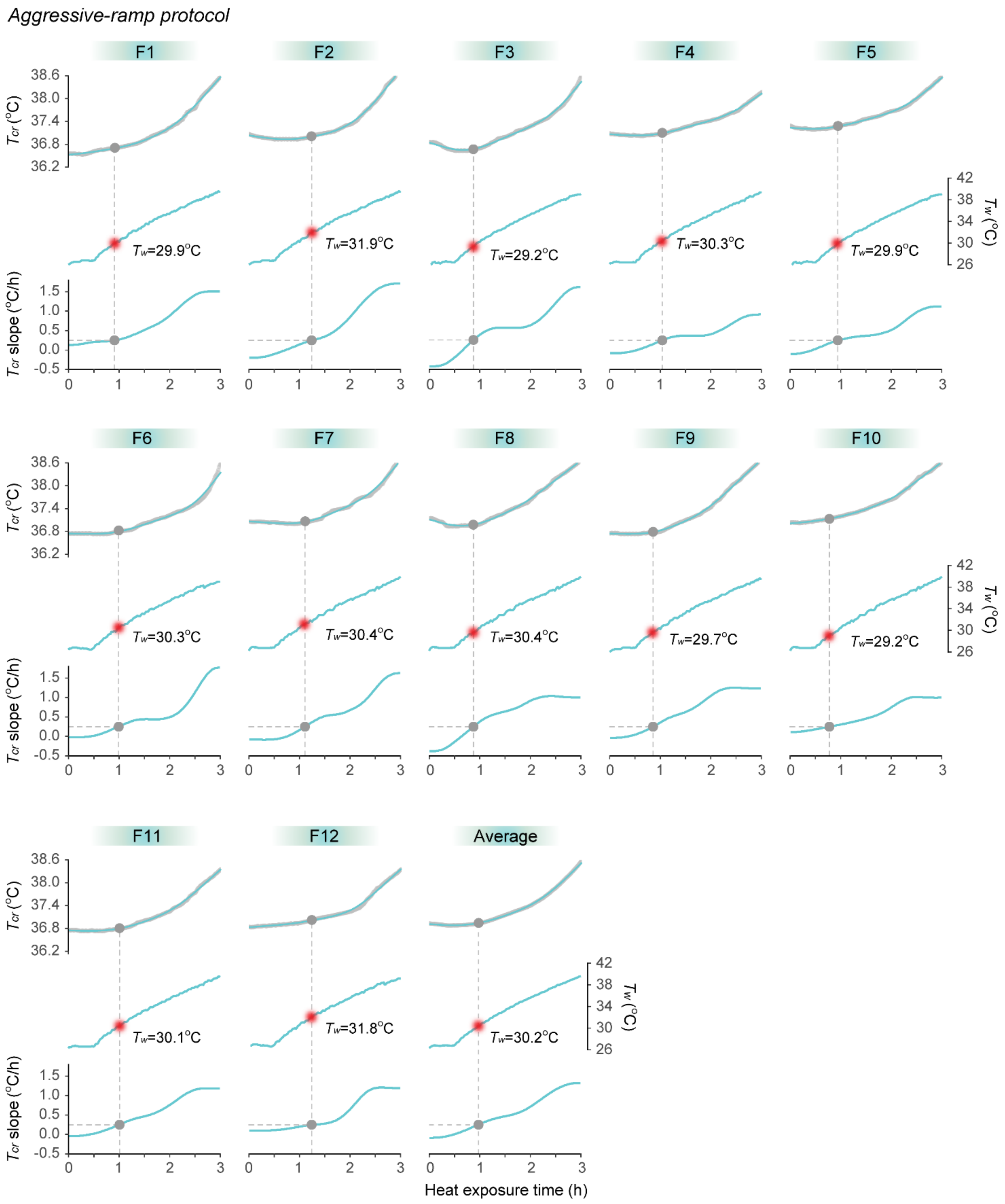


**Figure S3** Temporal variations in individual and average core temperature response, recorded environmental wet-bulb temperature in the heat chamber, and the slope of the rectal temperature changes under the aggressive-ramp protocol at a dry-bulb temperature of 42 °C in young adult females. F1–F12 denote female participants 1 through 12.

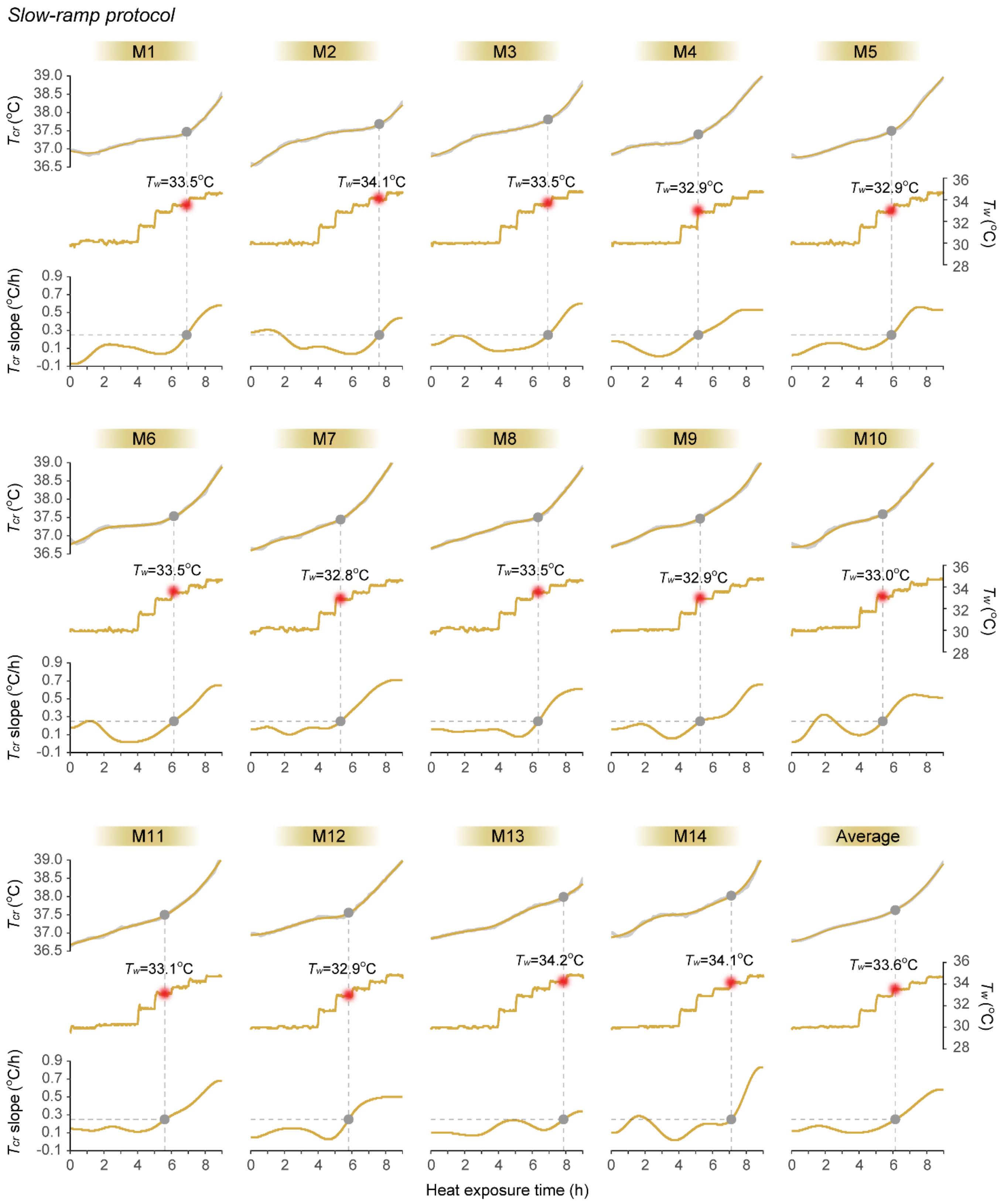


**Figure S4** Temporal variations in individual and average core temperature response, recorded environmental wet-bulb temperature in the heat chamber, and the slope of the rectal temperature changes under the slow-ramp protocol at a dry-bulb temperature of 42 °C in young adult males. M1–M14 denote male participants 1 through 14.

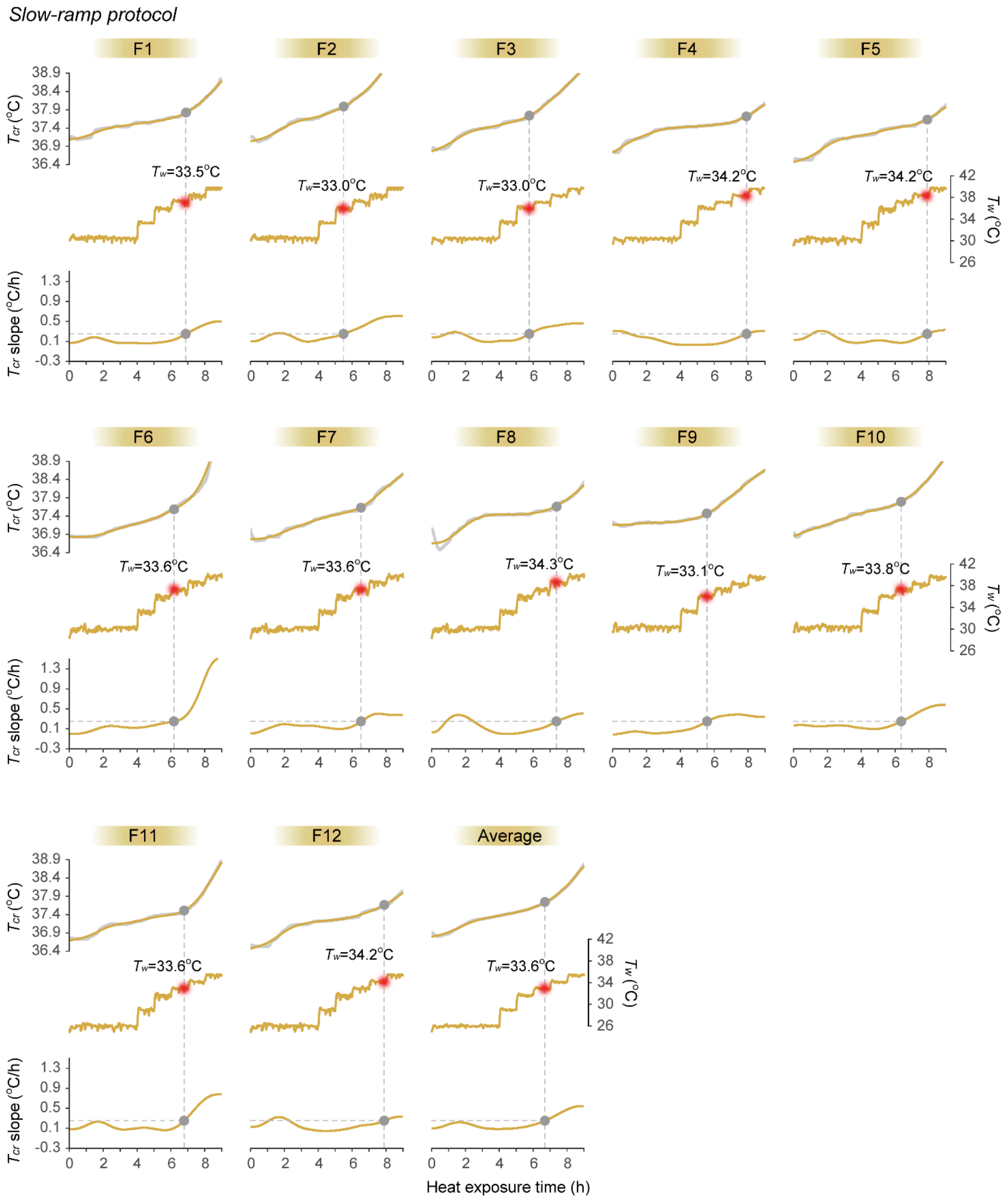


**Figure S5** Temporal variations in individual and average core temperature response, recorded environmental wet-bulb temperature in the heat chamber, and the slope of the rectal temperature changes under the slow-ramp protocol at a dry-bulb temperature of 42 °C in young adult females. F1–F12 denote female participants 1 through 12.

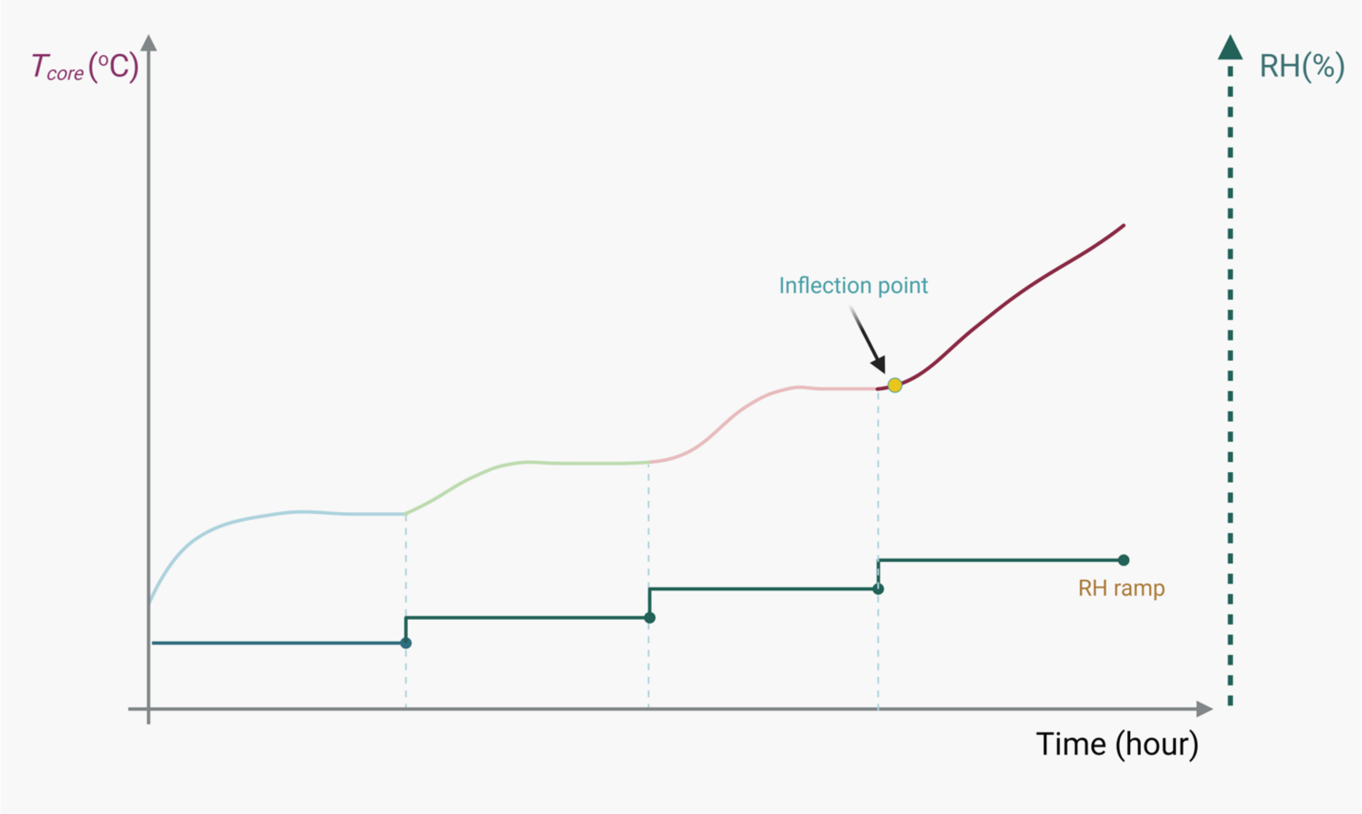


**Figure S6** Schematic illustration of an ideal humidity ramp protocol. Each stepwise increase in relative humidity (RH) is maintained for a sufficiently long duration to allow core temperature stabilization before progressing to the subsequent RH level.

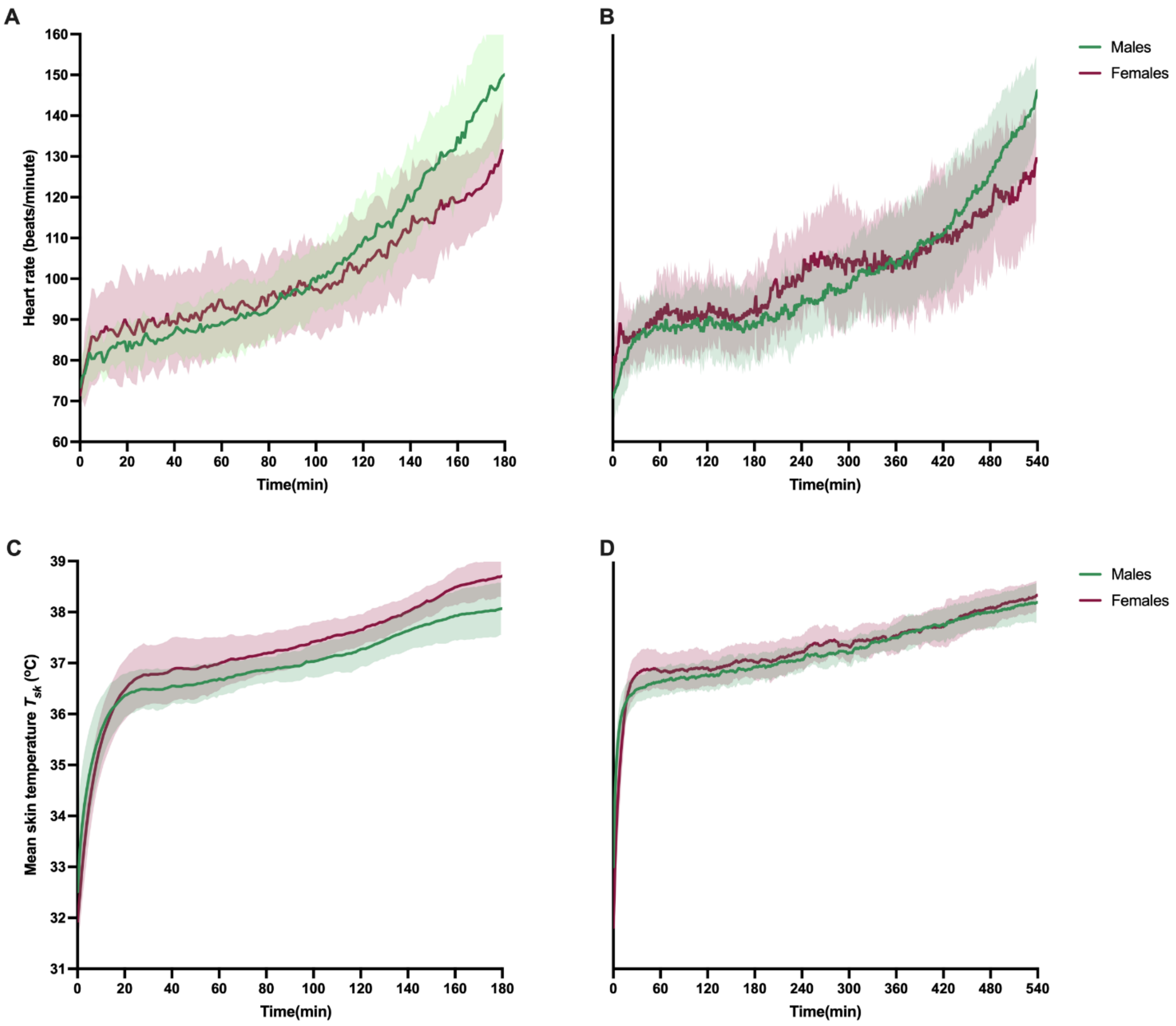


**Figure S7** Temporal changes in average heart rate and skin temperature responses during aggressive-ramp and slow-ramp humidity protocols at a fixed dry-bulb temperature of 42°C in males and females. **A**, heart rate progression under the aggressive-ramp protocol; **B**, heart rate progression under the slow-ramp protocol; **C**, mean skin temperature responses under the aggressive-ramp protocol; **D**, mean skin temperature responses under the slow-ramp protocol.

### Supplemental tables

**Table S1** Sensitivity analysis of critical inflection point (CIP) determination using alternative smoothed core temperature slope thresholds (0.15, 0.20, 0.25, and 0.30 °C/h). The analysis assessed the robustness of CIP identification for the compensable–uncompensable heat stress transition. The results indicate that the 0.25 °C/h threshold yielded stable and physiologically consistent estimates, whereas less stringent or more stringent thresholds advanced or delayed the transition unrealistically, potentially obscuring the early onset of uncompensable heat stress. "NA" denotes that a critical infection point could not be identified at the specific threshold because a predefined termination criterion was reached prior to CIP detection.

| | CIPs derived from the aggressive-ramp protocol (°C) | | | | CIPs from the slow-ramp protocol (°C) | | | |
|---|---|---|---|---|---|---|---|---|
| Slope threshold | 0.15°C/h | 0.20°C/h | 0.25°C/h | 0.30°C/h | 0.15°C/h | 0.20°C/h | 0.25°C/h | 0.30°C/h |
| M1 | 27.8 | 27.8 | 27.8 | 32.7 | 33.5 | 33.5 | 33.5 | 34.0 |
| M2 | 27.8 | 27.8 | 27.8 | 27.8 | 34.2 | 34.2 | 34.1 | 33.9 |
| M3 | 27.8 | 27.8 | 31.8 | 32.9 | 33.5 | 33.6 | 33.5 | 34.2 |
| M4 | 27.7 | 27.7 | 31.8 | 33.2 | 31.5 | 31.6 | 32.9 | 32.9 |
| M5 | 27.7 | 27.7 | 29.0 | 29.3 | 32.9 | 32.9 | 32.9 | 33.5 |
| M6 | 27.7 | 27.7 | 29.4 | 33.5 | 32.8 | 32.8 | 33.5 | 33.5 |
| M7 | 27.9 | 27.9 | 29.8 | 31.0 | 30.3 | 31.4 | 32.8 | 32.9 |
| M8 | 27.9 | 27.9 | 27.9 | 27.9 | 32.9 | 33.6 | 33.5 | 33.4 |
| M9 | 31.1 | 31.5 | 31.9 | 32.3 | 31.5 | 31.5 | 31.6 | 33.5 |
| M10 | 29.5 | 30.6 | 31.4 | 31.8 | 31.7 | 33.3 | 33.0 | 33.1 |
| M11 | 29.0 | 29.5 | 29.9 | 30.5 | 31.7 | 33.1 | 33.1 | 33.6 |
| M12 | 28.9 | 29.4 | 31.6 | 32.4 | 32.9 | 32.9 | 32.9 | 32.8 |
| M13 | 28.1 | 28.1 | 28.1 | 28.1 | 31.0 | 31.6 | 34.2 | NA |
| M14 | 30.1 | 30.2 | 30.6 | 32.0 | 31.5 | 33.6 | 34.1 | 34.1 |
| F1 | 26.8 | 26.8 | 29.9 | 30.9 | 33.7 | 33.7 | 33.5 | 34.3 |
| F2 | 30.2 | 31.0 | 31.9 | 32.6 | 31.8 | 31.6 | 33.0 | 33.0 |
| F3 | 28.8 | 29.0 | 29.2 | 29.5 | 33.1 | 33.0 | 33.0 | 33.4 |
| F4 | 29.2 | 29.8 | 30.3 | 31.0 | 34.3 | 34.2 | 34.2 | NA |
| F5 | 28.3 | 29.0 | 29.9 | 30.9 | 34.3 | 34.2 | 34.2 | NA |
| F6 | 29.2 | 29.7 | 30.3 | 30.9 | 31.7 | 33.0 | 33.6 | 33.5 |
| F7 | 30.4 | 30.8 | 30.4 | 31.3 | 33.2 | 33.6 | 33.6 | 33.6 |
| F8 | 29.2 | 29.3 | 30.4 | 29.8 | 33.7 | 34.5 | 34.3 | 34.3 |
| F9 | 28.6 | 29.1 | 29.7 | 29.9 | 31.7 | 33.0 | 33.1 | 32.9 |
| F10 | 26.8 | 27.3 | 29.2 | 30.2 | 32.9 | 33.2 | 33.8 | 33.8 |
| F11 | 28.9 | 29.5 | 30.1 | 30.7 | 33.6 | 33.7 | 33.6 | 33.6 |
| F12 | 28.7 | 30.3 | 31.8 | 33.5 | 33.6 | 34.2 | 34.2 | NA |

**Note**: M1–M14 denote male participants 1 through 14, and F1–F2 denote female participants 1 through 12.

**Table S2** Urine specific gravity, total sweat production, sweating rate, and perceptual responses (thermal sensation, thermal discomfort, thirst sensation, and skin wetness sensation) in young adult males ($n$ = 14) and young adult females ($n$ = 12) during exposure to $T_{db}$=42 °C under two humidity ramp protocols (aggressive ramp and slow ramp). Paired-samples $t$-tests were used to compare pre-exposure values and post-exposure values with each protocol (aggressive ramp and slow ramp) as well as corresponding values between the aggressive-ramp and slow-ramp conditions. Sex differences were assessed using independent $t$-tests with Welch's correction for unequal variances. *, indicates a significant difference between pre-exposure and post-exposure ($p < 0.05$); $^{\#\#}$, indicate a significant difference between sexes ($p < 0.001$); -, not applicable.

| **Indicators** | **Time point** | **Aggressive-ramp** | | **Slow-ramp** | |
|---|---|---|---|---|---|
| | | Males | Females | Males | Females |
| Urine specific gravity | Pre-exposure | 1.013±0.006 | 1.013±0.007 | 1.010±0.006 | 1.013±0.007 |
| | Post-exposure | 1.014±0.004 | 1.013±0.007 | 1.010±0.006 | 1.011±0.008 |
| Total sweat production (g) | - | 1091±341 | 1116±376 | 2981±629 | 3273±887$^{\#\#}$ |
| Total exposure duration (min) | - | 153.8±14.2 | 159.1±8.3 | 488.9±41.9 | 498.3±34.9 |
| Sweating rate (g/h) | - | 425.6±112.6 | 422.3±145.7 | 368.8±72.1 | 390.1±83.0 |
| Dehydration status | - | Euhydrated | Euhydrated | Euhydrated | Euhydrated |
| Metabolic rate (METs) | Pre-exposure | 1.39±0.13 | 1.28±0.11 | 1.36±0.15 | 1.25±0.12 |
| | Post-exposure | 1.48±0.15 | 1.38±0.09 | 1.46±0.14 | 1.36±0.11 |
| **Indicators** | **Time point** | **Perceptual responses** | | | |
| | | **Males** | **Females** | **Males** | **Females** |
| Thermal sensation | Pre-exposure | -0.07±0.47 | -0.08±0.29 | -0.07±0.27 | -0.17±0.39 |
| | Post-exposure | +2.43±0.65 | +2.83±0.39 | +2.78±0.58 | +2.83±0.39 |
| Thermal discomfort | Pre-exposure | +0.15±0.38 | +0.17±0.39 | +0.38±0.65 | +0.17±0.39 |
| | Post-exposure | -2.54±0.97 | -2.83±0.39 | -2.69±0.75 | -2.42±0.79 |
| Thirst sensation | Pre-exposure | 1.00±0.00 | 1.00±0.00 | 1.00±0.00 | 1.00±0.00 |
| | Post-exposure | 1.71±0.83 | 1.42±0.67 | 1.43±0.51 | 1.58±0.90 |
| Skin wetness sensation | Pre-exposure | 0.00±0.00 | 0.00±0.00 | 0.00±0.00 | -0.08±0.29 |
| | Post-exposure | -2.43±0.94 | -2.42±0.67 | -2.79±0.70* | -2.58±0.67 |
